# Prediction of virus-host associations using protein language models and multiple instance learning

**DOI:** 10.1101/2023.04.07.536023

**Authors:** Dan Liu, Francesca Young, David L Robertson, Ke Yuan

## Abstract

Predicting virus-host associations is essential to determine the specific host species that viruses interact with, and discover if new viruses infect humans and animals. Currently, the host of the majority of viruses is unknown, particularly in microbiomes. To address this challenge, we introduce EvoMIL, a deep learning method that predicts the host species for viruses from viral sequences only. It also identifies important viral proteins that significantly contribute to host prediction. The method combines a pre-trained large protein language model (ESM) and attention-based multiple instance learning to allow protein-orientated predictions. Our results show that protein embeddings capture stronger predictive signals than sequence composition features, including amino acids, physiochemical properties, and DNA k-mers. In multi-host prediction tasks, EvoMIL achieves median F1 score improvements of 8.6%, 12.3%, and 4.1% in prokaryotic hosts, and 0.5%, 1.8% and 3% in eukaryotic hosts. EvoMIL binary classifiers achieve impressive AUC over 0.95 for all prokaryotic and range from roughly 0.8 to 0.9 for eukaryotic hosts. Furthermore, EvoMIL estimates the importance of single proteins in the prediction task and maps them to an embedding landscape of all viral proteins, where proteins with similar functions are distinctly clustered together, highlighting the ability of EvoMIL to capture key proteins in virus-host specificity.

**Author summary:** Being able to predict which viruses can infect which host species, and identifying the specific proteins that are involved in these interactions, are fundamental tasks in virology. Traditional methods for predicting these interactions rely on common manual features among proteins, overlooking the structure of the protein ”language” encoded in individual proteins. We have developed a novel method that combines a protein language model and multiple instance learning to allow host prediction directly from protein sequences, without the need to extract manual features. This method significantly improved prediction accuracy and revealed key proteins involved in virus-host interactions.

## 1 Introduction

Advances in sequencing technologies, particularly metagenomics, have resulted in the identification of many new viruses. However, more than 90% of the virus sequences held in publicly available databases are not annotated with any host information [1].

Currently, there are no high-throughput experimental methods that can definitively assign a host to these uncultivated viruses. With a growing number of viruses being discovered, relying only on experiments to identify virus-host associations is challenging.

A number of computational approaches have been developed to predict unknown virus-host species associations. The coevolution of a virus and its host leave signals in virus genomes arising from the virus-host interaction. These signals have been exploited for in silico prediction of virus-host associations from virus genomes alone and fall into two broad types: 1) alignment-based approaches that search for homology such as prophage [2], CRISPR-cas spacers [3, 4] of genes acquired from [5]; 2) Alignment-free methods that use features such as k-mer composition, codon usage, or CpG content to measure the similarity between viral and host sequences or to other viruses with a known host[6]. To date, no computational approaches consider the structure of proteins from viruses for host species prediction purposes.

Here, we present a virus-host prediction model combining protein language models (PLM) and multiple instance learning (MIL). Transformers are self-supervised deep learning models [7] that learn the relationships among words within a sentence, and now dominate the field of natural language processing. More recently, the same architecture has been applied in biology, where words are replaced by amino acids and sentences by protein sequences. These transformer-based protein language models generate protein embeddings that encode structural features inferred from amino acid sequences based on large-scale protein databases [8]. Protein language models are trained on publicly available protein sequence archives and learn multi-level biological information from physiochemical properties of the individual amino acids to structural and functional information about proteins. Multiple instance learning (MIL) is a form of supervised learning that was developed for image processing tasks [9]. Instead of using individually labelled instances for classification, multiple instances are arranged together in a bag with a single label and classified together. We use attention-based MIL [10], which has the additional advantage of weighting instances in a bag, thereby indicating the importance of each instance in prediction.

The combination of the two approaches is particularly suited for virus-host prediction, as virus proteins collectively contribute to the association with a host. Instead of relying on predefined features, protein language models provide automatically learned features, free from design biases and limitations of the previous approaches. The ability to measure similarity and differences between protein sequences further boosts prediction performance through multiple instance learning, where viral proteins enabling interaction with hosts are highlighted through unbiased weight estimation.

In this paper, we introduce EvoMIL a method for predicting virus-host associations by combining the (Evo)lutionary Scale Modeling with (M)multiple (I)instance (L)earning, Fig 1. EvoMIL uses the model ESM-1b [8] to transform viral protein sequences into embeddings (i.e. numerical vectors) that are then used as features for virus-host classification. Multiple-instance learning allows us to consider each virus as a “bag” of proteins. We demonstrate that the embeddings capture the host signal from the viral sequences achieving high prediction scores at the species level of both prokaryotic and eukaryotic hosts. Furthermore, attention-based MIL enables us to identify which proteins are highly important in driving prediction and by implication are important to virus-host specificity.

**Fig 1.**
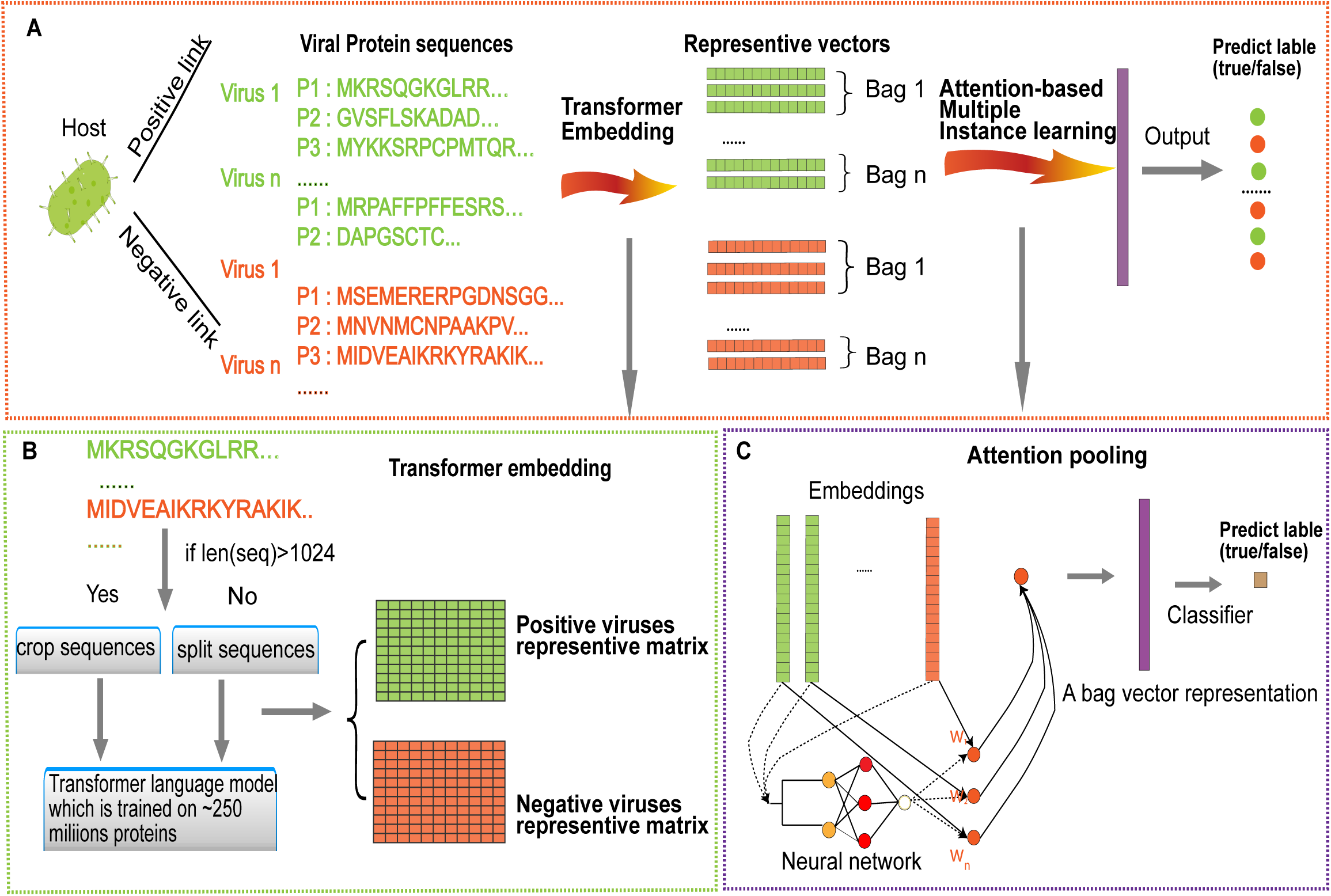
A diagrammatic representation of the EvoMIL method. A. Protein sequences of viruses and virus-host associations are collected from the VHDB [11]. For each host, we collect the same number of positive and negative viruses, and then embeddings of protein sequences from viruses are obtained by the pre-trained transformer model [8], which are features for host predictions based on attention-based MIL; B. Protein sequences of viruses are split to sub-sequences, which are used as input to the pre-trained transformer model to obtain the corresponding embeddings; C. There is a host label for a set of protein sequences on each virus, and attention-based MIL is applied to train the model for each host dataset by protein embeddings of viruses. Finally, we predict the host label for each virus and assign an instance weight that represents the importance of each protein for the virus.

## 2 Results

### 2.1 A benchmarking dataset for predicting virus-host association

Balanced binary datasets were generated from known virus-host associations documented in the Virus-Host database (VHDB) [11] for all hosts with a minimum threshold number of associations. These datasets consist of either all prokaryotic or all eukaryotic viruses. ’Positive’ viruses are those that are reported to be associated with the given host species. A matching number of ’negative’ viruses are randomly sampled from all other prokaryotic or eukaryotic viruses. The prokaryote datasets consist of nearly all dsDNA viruses which have 45 to 212 proteins coded in their genomes (Table S1), while the eukaryotic datasets include many RNA viruses that contain fewer proteins, ranging from 2 to 23 protein sequences (Table S2). The performance of MIL improves with higher numbers of instances in each bag, therefore we need to increase the threshold for the number of viruses in the eukaryotic training datasets to achieve similar performance with MIL. Accordingly, we set a threshold for minimum positive dataset size to 50 and 125 viruses for constructing prokaryotic and eukaryotic binary datasets, respectively. The aim of setting the threshold is to generate a sufficient number of validation samples for prokaryotic and eukaryotic hosts, respectively. Finally, we generated 15 prokaryotic host datasets and 5 eukaryotic host datasets.

For training the binary classification models, we use two strategies to sample the negative viruses: **Strategy 1** is to subsample viruses from those that infect any host from a different genus than those infected by the positive viruses; this means that the negative samples will be more distinct from the positive samples. **Strategy 2** is to select from those viruses that are associated with hosts belonging to the same taxonomic groups at the ranks of genus, family, order, class, and phylum as those hosts infected by positive viruses. In this way, the negative samples and positive samples are more likely to share structural mimics[12], so it will be challenging to train classifier models and make the binary models sensitive to capture the difference between positive and negative samples. The number of viruses related to each host is shown in Table S1. The largest prokaryote dataset is Mycolicibacterium smegmatis with 838 known viruses, followed by Escherichia coli with approximately half the number of viruses. For the eukaryotic datasets, Homo sapiens have by far the largest number of known virus species (1321) with the next highest being the tomato (Solanum lycopersicum) at 277, see Table S2.

### 2.2 EvoMIL achieves high performance for binary virus-host prediction

Embedding vectors for each of the proteins of a virus generated with the protein language model, ESM1b, were used as an instance in a “virus bag” for MIL. These labelled bags were used to train the MIL model using 5-fold cross-validation on 80% of the datasets, then each 5-fold model performance was evaluated on the remaining 20% of the datasets. Each model is evaluated with a range of metrics: AUC, accuracy, F1 score, sensitivity, specificity, and precision. We evaluated the predictive performance of EvoMIL for binary classification using the datasets generated with both **Strategy 1** and **Strategy 2** above, training a prediction model for each host.

#### 2.2.1 Prokaryotic and Eukaryotic host performance

The heatmaps of evaluation indices for the prokaryotic and eukaryotic host classifiers are presented in Fig 2 A and C. Here, evaluation indices are calculated based on the best-performing model with the highest AUC chosen in 5-fold cross-validation. In Fig 2 A, the accuracy is higher than 0.9 except for two hosts, which are 0.86. The ROC curves in Fig 2 B show that all prokaryotic classifiers perform very strongly with each host achieving an AUC greater than 0.95 and 10 achieving an AUC of 1.

**Fig 2.**
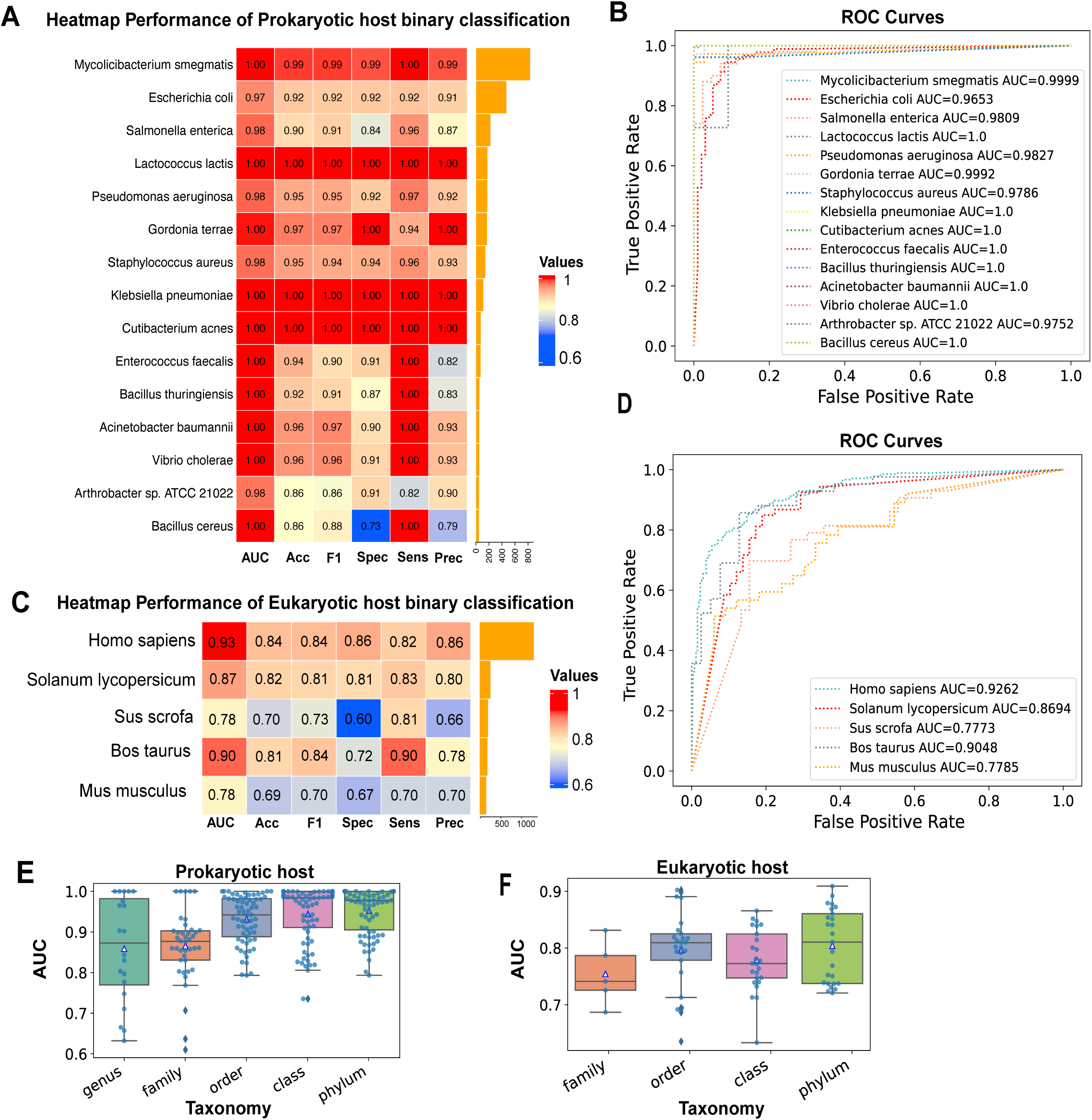
Performance of binary classification tasks. This figure separately shows the heatmap of AUC, accuracy, F1 score, sensitivity, specificity, and precision on 15 prokaryotic (A) and 5 eukaryotic host binary classifiers (C), negative samples are selected by **strategy 1**; ROC curves of 15 prokaryotic hosts (B) and 5 eukaryotic hosts (D) corresponding with heatmap plots A and C; AUC of different taxonomy on prokaryotic (E) and eukaryotic hosts (F) where negative samples are selected using **strategy 2**.

We also obtained the mean and standard deviation of each host, by testing 5-fold cross-validation models on the host test dataset (see Table S3). EvoMIL shows good performance, with 14/15 hosts achieving a mean AUC greater than 0.9. Overall, our results demonstrated that EvoMIL shows an impressive performance in the binary classification tasks of viruses associated with prokaryotic hosts. More evaluation metrics are included in Table S3.

The accuracy of each eukaryotic host classifier is shown in Fig 2 C, it is clear that all hosts perform with an accuracy higher than 0.8 except for two hosts which are roughly 0.7. H. sapiens obtained the highest accuracy with 0.84; Mus musculus has the lowest accuracy with 0.69. The ROC curves of the 5 eukaryotic hosts classifiers are presented in Fig 2 D. Although the eukaryote classifiers achieve good performance with AUCs above 0.77, they perform less well than the prokaryote classifiers, with only 3/5 datasets scoring an AUC above 0.85. There may be several explanations for the lower performance. Firstly, the average number of proteins per virus is much lower, resulting in small “bags” for MIL. Secondly, there is a much higher diversity of virus types in the datasets of the eukaryotic hosts, often containing viruses from multiple Baltimore classes. The virus of these different classes is polyphyletic meaning they will have no common ancestor and therefore have no shared genes and interact with different host pathways.

The mean and standard deviation of each host are obtained by testing five trained cross-validation models on the test data set (see Table S4). Here, the mean AUC is higher than 0.85 except for the classifiers of two hosts that perform less well, with AUC scores of 0.761*±*0.01 for Sus scrofa and 0.762*±*0.02 for M. musculus. Overall, our results demonstrate that EvoMIL performs well in binary classification tasks of viruses associated with eukaryotic hosts.

#### 2.2.2 Sampling negative samples from similar viruses makes binary host classification more challenging

Next, we test our model with more challenging tasks. Using the second strategy of selecting negative viruses that are associated with hosts sharing the same taxonomic rankings as the hosts associated with positive viruses, we observe that the classification task becomes increasingly challenging as we move from the phylum to the genus level. Results show that our EvoMIL models achieve high AUC scores but that distinguishing between viruses of similar hosts is more difficult with a noticeable drop in performance at family and genus levels. In Fig 2, the box plots show AUC of prokaryotic (E) and eukaryotic (F) hosts based on negative selection **strategy 2** with five taxonomies genus/family/order/class/phylum. Phylum level (lime colour) presented significantly improves compared with lower taxonomies, especially genus level. Note, at the lower taxonomic ranks there are only sufficient numbers of negative viruses to meet our threshold of 50 for 4 hosts at the genus level, 8 hosts at the family level and 13 hosts at the order level.

### 2.3 Embedding features outperform protein and DNA k-mer features on Multi-class tasks

To demonstrate that the ESM embeddings are encoding highly predictive information than conventional features, we generated feature sets from k-mer composition of the nucleic acid, amino-acid and physio-chemical sequences [13], and implemented ESM-1b and k-mer features on multi-class classification model. Here, we performed multi-class classification extending attention-based MIL by modelling the joint Multinomial distribution of bag labels. Both prokaryotic and eukaryotic multi-class datasets were constructed using hosts infected by at least 30 viruses. This resulted in 22 classes for the prokaryotes and 36 classes for the eukaryotes. Again, we applied 5-fold cross-validation on training datasets, then separately tested trained models on the testing dataset. These models are named ESM-1b, AA 2 and PC 3 and DNA 5, according to different features.

The results for each model are presented in Table 1, we obtain AUC, accuracy, and F1 scores by evaluating each of the 5-fold cross-validation models with the test dataset. Comparing ESM-1b with AA 2, PC 4, and DNA 5 for both prokaryotic and eukaryotic hosts, the ESM-1b features have the strongest prediction signal since during testing on prokaryotic hosts, the mean F1 score of EvoMIL is 0.88 which is 8.6%, 12.3%, and 4.1% higher compared with AA 2, PC 4 and DNA 5, respectively; while the counterpart of EvoMIL in eukaryotic hosts is 0.292 which is 0.5%, 1.8% and 3% higher compared with AA 2, PC 4 and DNA 5 (Table 1). Additionally, ESM-1b demonstrates better performance in terms of AUC and accuracy compared to k-mers, except for eukaryotes, the accuracy of ESM-1b is comparable to that of AA 2.

**Table 1.** The AUC, Accuracy and F1 score of multi-class MIL by using ESM-1b and k-mer features Here is a comparison of AUC, accuracy, F1 score between ESM-1b and k-mer feature sets(AA 2, PC 4 and DNA 5). For each feature, the evaluation was conducted by training multi-class classification models on 22 prokaryotic hosts and 36 eukaryotic hosts, and the mean and standard deviation of the evaluation metrics (AUC, accuracy, and F1 score) were obtained using 5-fold cross-validation.

| Host type | Methods | AUC | Accuracy | F1 score |
| --- | --- | --- | --- | --- |
| **Prokaryotes** | ESM-1b | **0.992±0.0** | **0.909±0.0** | **0.88±0.01** |
|  | AA_2 | 0.979±0.0 | 0.856±0.01 | 0.794±0.02 |
|  | PC_3 | 0.969±0.01 | 0.843±0.01 | 0.757±0.02 |
|  | DNA_5 | 0.987±0.0 | 0.882±0.01 | 0.839±0.01 |
| **Eukaryotes** | ESM-1b | **0.831±0.01** | **0.494±0.01** | **0.292±0.01** |
|  | AA_2 | 0.829±0.03 | 0.494±0.01 | 0.287±0.01 |
|  | PC_3 | 0.821±0.01 | 0.479±0.01 | 0.274±0.02 |
|  | DNA_5 | 0.801±0.02 | 0.466±0.02 | 0.262±0.01 |

Fig 3 A and C, respectively, show AUC and accuracy across ESM-1b and k-mer features in both prokaryotes and eukaryotes, and AUC and accuracy are equivalent with those presented in Table 1. In Fig 3 A, ESM-1b performs the highest AUC and smallest standard deviation among prokaryotic hosts; for eukaryotic hosts, ESM-1b shows the smallest standard deviation and the highest mean AUC compared with k-mer features. Here, AA 2 has the largest variation, despite obtaining the highest AUC. In Fig 3 C, ESM-1b presents the highest accuracy and the smallest standard deviation among prokaryotic hosts; for eukaryotic hosts, ESM-1b includes the highest accuracy.

**Fig 3.**
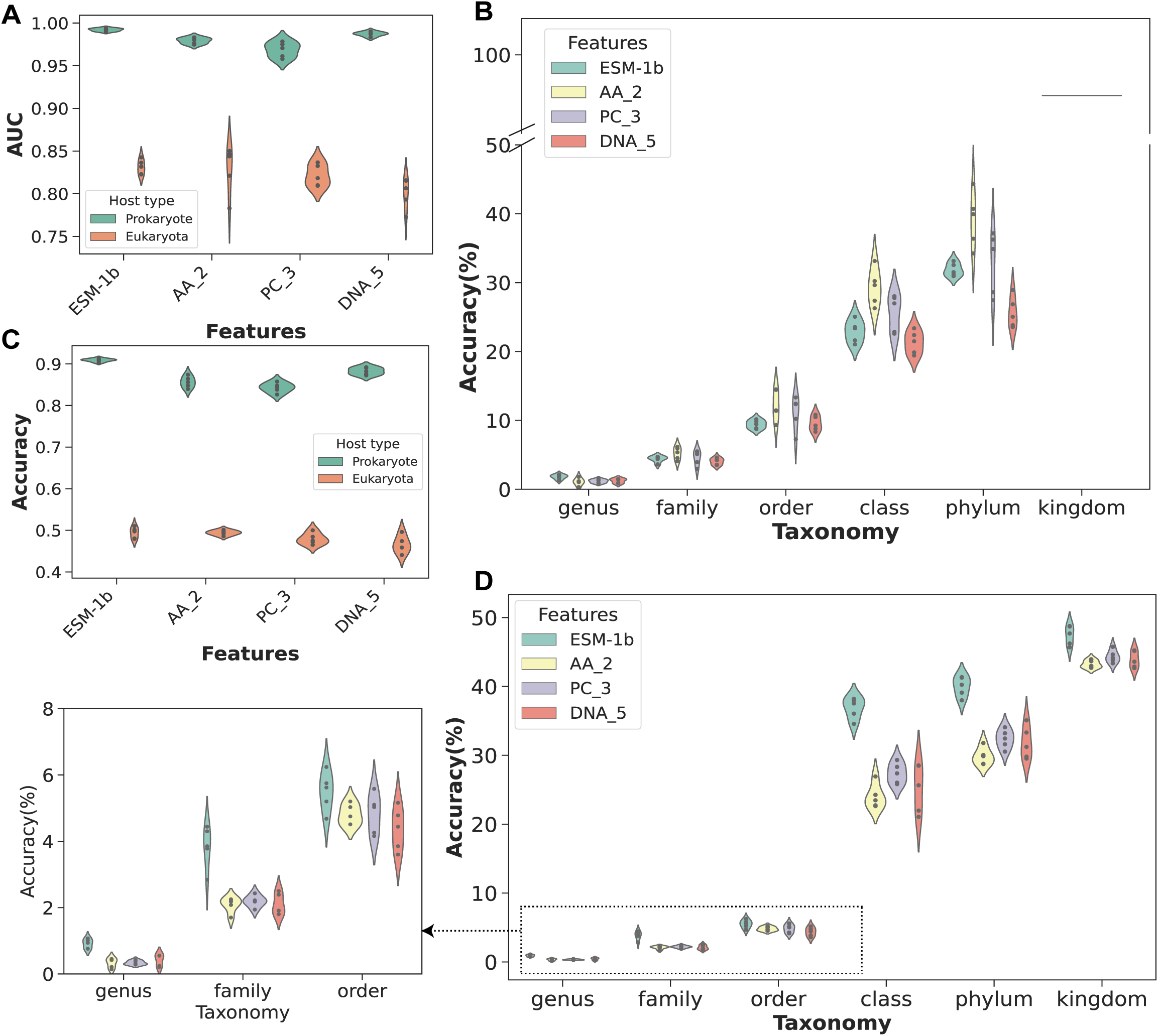
A and C represent the AUC and accuracy, respectively, for prokaryotic and eukaryotic hosts using four feature sets (ESM-1b, AA 2, PC 3 and DNA 5). B and D indicate the results obtained by testing the trained models on prokaryotic and eukaryotic hosts associated with 5 to 30 viruses, using the four different feature sets.

Additionally, although AA 2 presents the smallest standard deviation in accuracy, ESM-1b obtained the best mean AUC and F1 score (see Table 1).

To evaluate the multi-class MIL models, we select those hosts associated with fewer viruses, ranging from 5 to 30. ESM-1b and k-mer features (AA 2, PC 4, and DNA 5) can be extracted for these viruses, then they can be tested based on trained multi-class host models. Based on 5-fold cross-validation, Fig 3 B and D illustrate the accuracy of ESM-1b and k-mer features on MIL multi-class models. Here, if predicted host labels belong to the same genus/family/order/class/phylum/kingdom as the true host label, we consider it a correct prediction. Then we can calculate accuracy by determining the percentage of correct predictions on all test samples to evaluate the performance of EMS-1b and k-mer features.

In prokaryotic hosts (Fig 3 B), accuracy is roughly between 1% and 10% on genus, family and order levels, while accuracy is between 20% and 90% on class, phylum, and kingdom levels. It indicates that prediction is more challenging in lower taxonomies. ESM-1b performs best on the genus level with the highest mean accuracy, 1.8%, while the mean accuracy for AA 2, PC 3, DNA 5 are 1.09%, 1.18% and 1.23%, respectively (see Table S5). Furthermore, across each taxonomy level, the standard deviation of ESM-1b is the smallest compared with k-mer features, although ESM-1b does not perform with the highest accuracy on higher taxonomic levels. Overall, ESM-1b performs best on the genus level and shows stable accuracy results across all taxonomy levels on prokaryotic hosts.

As for eukaryotic hosts (Fig 3 D), accuracy is roughly between 1% and 6% on genus, family and order levels, while accuracy is between 25% and 50% on class, phylum, and kingdom levels. Across all taxonomic levels, ESM-1b consistently outperforms DNA and protein k-mer features in terms of accuracy. For example, the mean accuracy of ESM-1b on the class level is 36.868%, which is higher by 12.86%, 9.49%, and 11.72% than AA 2, PC 3 and DNA 5, respectively (see Table S5). In summary, the multi-class MIL model based on ESM-1b features shows potential in predicting hosts that are associated with fewer than 30 viruses in eukaryotic datasets.

Overall, ESM-1b demonstrated superior performance compared to the k-mer features, through 5-fold cross-validation on both prokaryotic and eukaryotic hosts. Furthermore, we split multi-class datasets into 80% training and 20% test sets, then train and test our model without cross-validation, and compare the accuracy for each host to evaluate the prediction performance of ESM-1b and k-mer features on multi-class classification.

Fig 4 A and B, respectively, presented taxonomy trees of prokaryotic and eukaryotic aligned with Log2 accuracy ratio between ESM-1b and K-mers (AA 2, PC 3 and DNA 5) for each host. Results show that ESM-1b achieved the highest accuracy in 17 out of 22 prokaryotic hosts compared with protein and DNA k-mer features. Similarly, ESM-1b outperforms the other feature sets in 16 out of 36 eukaryotes. These findings demonstrate ESM-1b performs best in both prokaryotic and eukaryotic host multi-classification tasks compared with k-mer feature sets.

**Fig 4.**
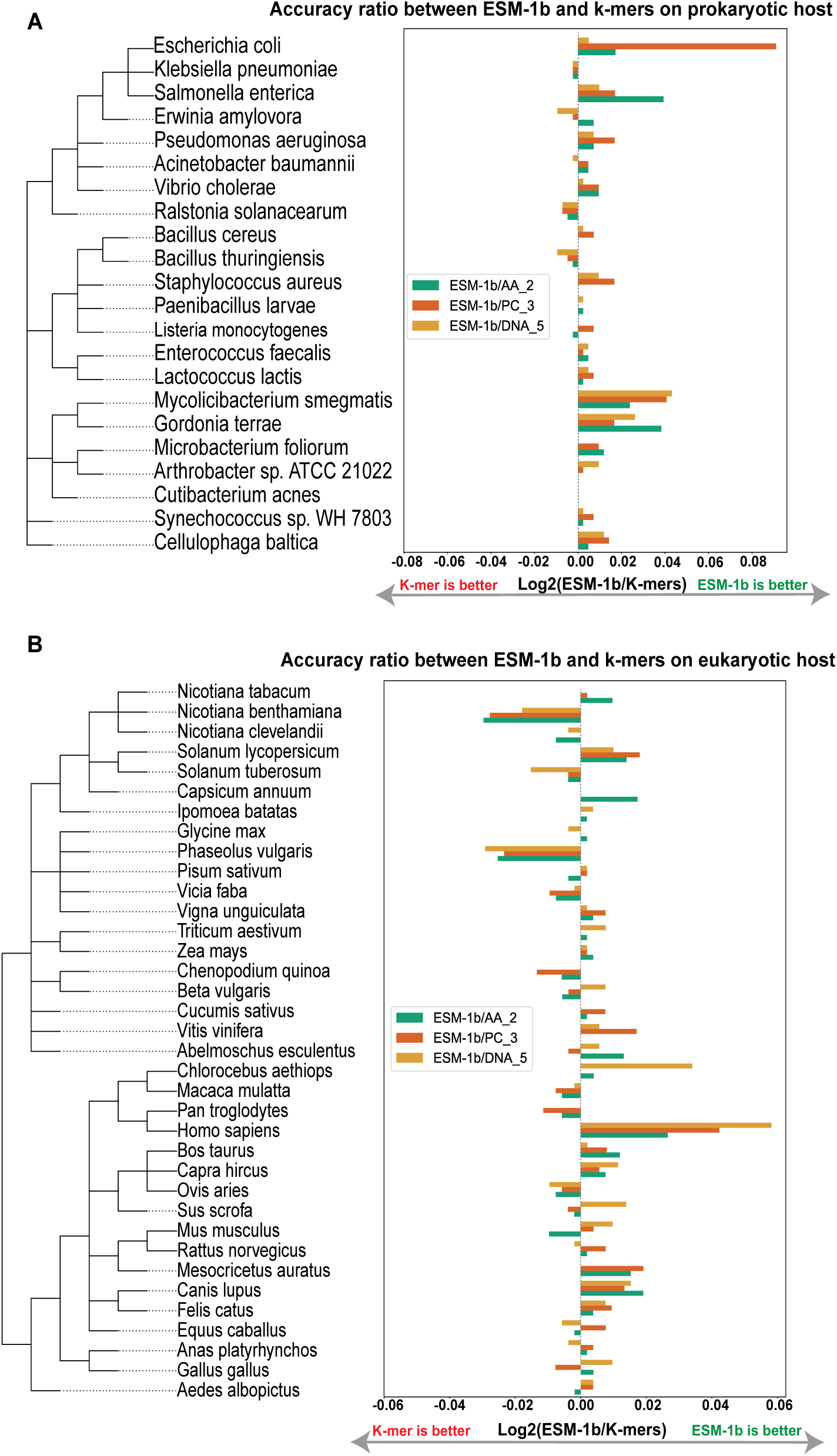
The taxonomy tree, aligning with Log2 of ratio accuracy between ESM-1b and k-mers. The figure shows the taxonomy tree of 22 prokaryotic (A) and 36 eukaryotic (B) hosts. Each host is aligned with a bar plot showing the accuracy ratio between ESM-1b and AA 2, PC 3, and DNA 5, respectively.

To understand if the host phylogeny explains the prediction performance of multi-class classification, we align the taxonomy tree of hosts with the heatmap showing the number of predicted false hosts for each host (Fig 5). The objective is to determine if false predicted host classes of viruses are more likely to share common parent nodes with true hosts in the taxonomy tree. In Fig 5, two heatmaps respectively present the number of false predicted hosts which belong to the same taxonomy as the true host in prokaryotic and eukaryotic hosts.

**Fig 5.**
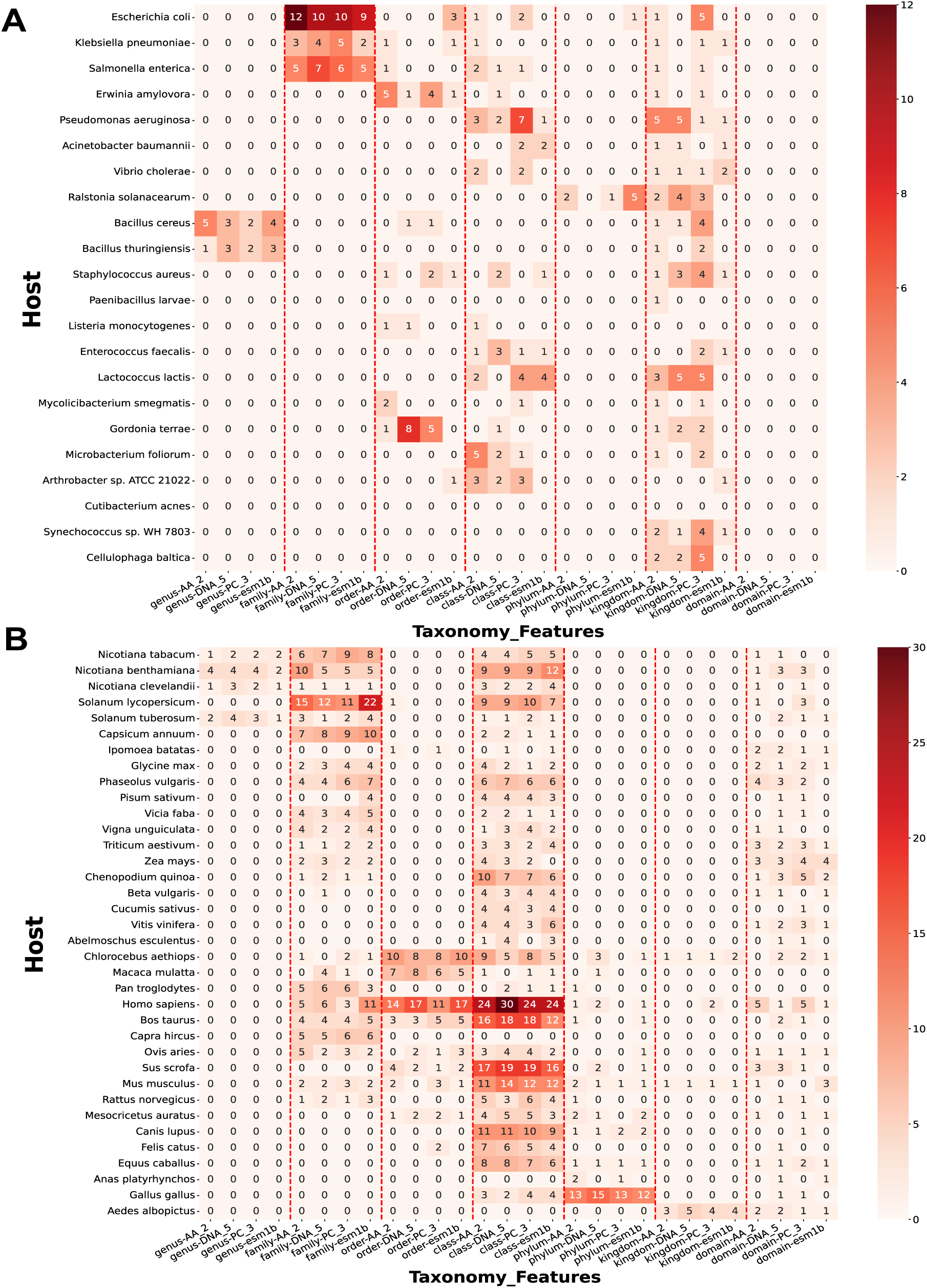
The figure represents heatmaps between different features and each prokaryotic (A) and eukaryotic (B) host. The values in the heatmap are the total number of predicted hosts which belong to the same taxonomy as the true host.

In the prokaryotic host heatmap (Fig 5 A), the highest number of falsely predicted hosts is observed at the family level. Notably, E. coli accounts for the largest number of false predicted hosts. Furthermore, an interesting observation is that Bacillus cereus and Bacillus thuringiensis host models predict each other as the host label (see Fig. S1), and they belong to the same genus, indicating that predicted hosts have a tendency to be hosts sharing the same taxonomy levels with true host label. In the eukaryotic host heatmap (Fig 5 B), the most number of false predicted hosts belong to the class level. H. sapiens and Bos taurus have the highest number of false predicted hosts, and they predict each other as the host label (see Fig. S2). Similarly, a similar situation can be observed between S. scrofa and M. musculus, which further reinforces the finding that closely related hosts within the taxonomic tree are more likely to be predicted as each other’s labels.

To understand the relationship between host phylogeny and predicted results, we use E. coli as an example in prokaryotes (see Fig 4 A). In Fig. S1, based on ESM-1b, the number of predicted false host labels are: 7 labels for S. enterica, 3 labels for Erwinia amylovora, 2 labels for K. pneumoniae, and 1 label for Ralstonia solanacearum, where S. enterica and K. pneumoniae belong to the same family as E. coli, E. amylovora belongs to the same order, and R. solanacearum belongs to the same phylum as E. coli. In Fig 4 B, it is clear that it is more challenging to predict eukaryotic hosts compared with prokaryotic hosts. ESM-1b demonstrates significantly superior performance compared to k-mer features in Nicotiana clevelandii and Cucumis sativus, while its performance on other hosts within the eukaryotic taxonomy tree is comparable to that of the k-mer features. In the case of H. sapiens as an example, the false positive host labels based on ESM-1b for H. sapiens primarily consist of 11 Pan troglodytes labels, 9 Chlorocebus aethiops labels, 8 Macaca mulatta labels, and 8 S. scrofa labels. Here, Pan troglodytes belongs to the same family as H. sapiens; Chlorocebus aethiops and Macaca mulatta belong to the same order level as H. sapiens; and S. scrofa belongs to the same class level as H. sapiens.

Overall, viruses are more likely to be misclassified as closely related hosts during multi-class classification, confirming that those hosts sharing common parent nodes in the taxonomy tree tend to be infected by similar viruses [13].

### 2.4 MIL attention weights can be used to interpret which virus proteins are important

Next, we investigate if the proteins identified as important for the prediction can be explained as proteins that are key contributors to virus-host specificity, i.e. whether the transformer embeddings are encoding biologically meaningful information about the viral proteins. As we described in the introduction, attention-based MIL learns the weight of each protein in a virus bag, where a high weight indicates that a protein is important for host prediction, and by implication is more likely to be important to virus-host specificity. We used UMAP[14] to investigate whether the embeddings of these important proteins identified by attention-based MIL contain any underlying clustering structures.

For this investigation, we used the viruses infecting E. coli and H. sapiens which contain the most annotations, because they are the most extensively studied hosts from each domain. Using the learned model parameters of the best-performing cross-validation model in the binary and multi-class classification tasks respectively, we ranked the attention weights of the proteins for the positive viruses and selected the top 5 ranked proteins for each virus. Furthermore, we collect two groups of top protein embeddings based on binary (Fig 6 A, C) and multi-class classification (Fig 6 B, D) models, respectively. These proteins were annotated with functions related to the viral life cycle using Gene Ontology (GO) terms obtained from InterProScan [15]. UMAPs were used to visualize any clustering structure that indicates similarities of the protein embeddings (see Fig 6).

**Fig 6.**
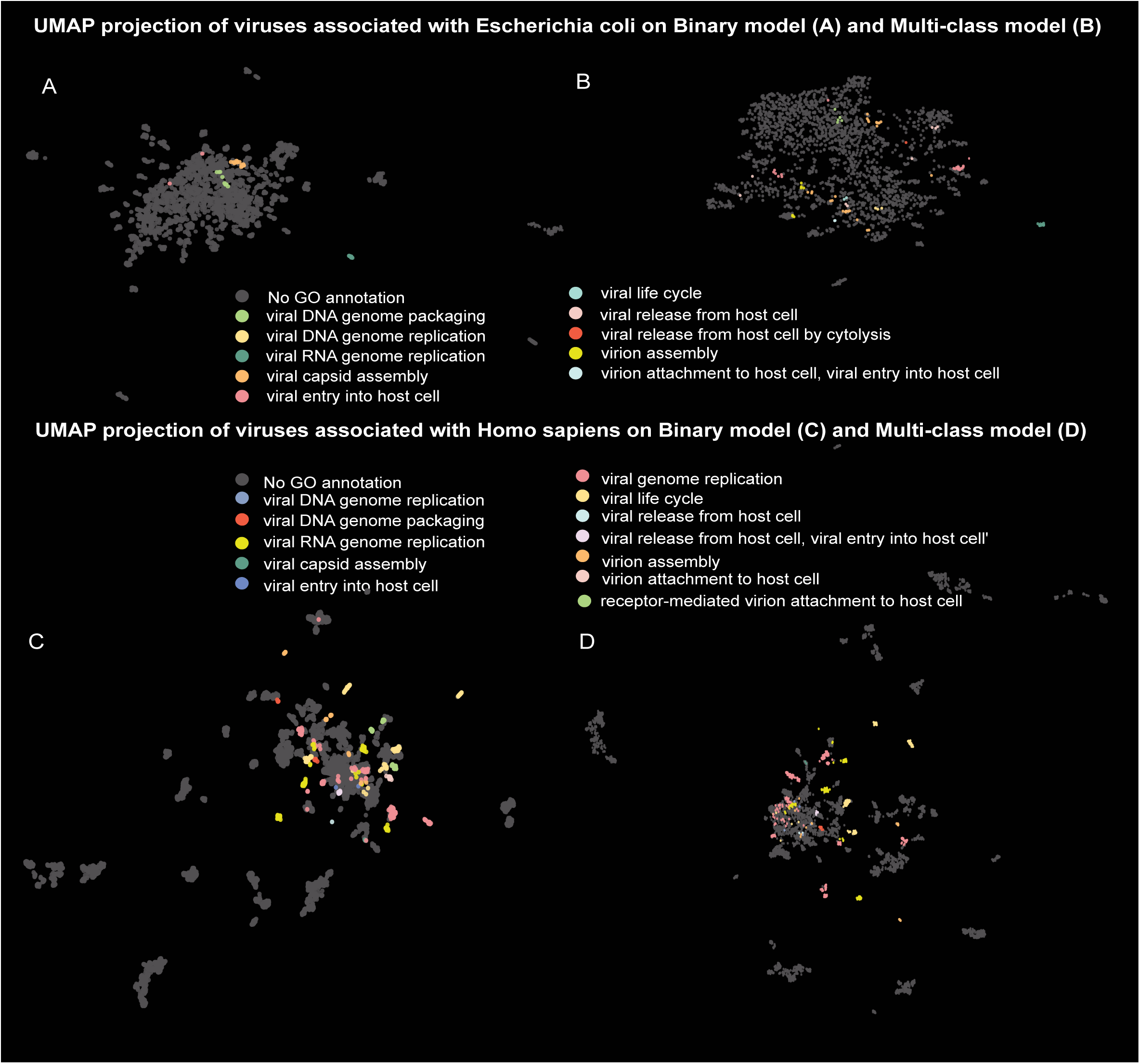
UMAPs projection of protein embeddings of viruses associated with E. coli (top) and H. sapiens (bottom). Figure A and B show the two-dimensional distribution of top 5 ranked protein embeddings for each virus associated with E. coli, while figure C and D show the distribution of embedding vectors of top 5 ranked proteins for each virus associated with H. sapiens. Here, the protein weights in A and C are obtained by binary models, whereas in B and D, the weights are obtained by multi-class classification models. Protein GO terms regarding the viral life cycle of viruses are highlighted with different colours, allowing us to understand predictive signals of proteins captured by the pre-trained transformer model.

Interestingly all UMAPs of E. coli and H. sapiens show distinct clustering structures, we assume that these clusters occur because the different functions driving host specificity are captured by the ESM-1b embeddings. In other words, ESM-1b embeddings encode functional and structural signals, and MIL learns some consistent features associated with host specificity.

For the set of top-ranked proteins of the viruses associated with E. coli based on the binary model, only roughly 1.5% (35 out of 2359) were assigned a viral life cycle GO term. In Fig 6 A, four different GO terms related to the viral life cycle were used to colour the proteins in the UMAP, illustrating that proteins with different GO terms form distinct clusters. More specificity, viral capsid assembly (GO:0019069), viral DNA genome packaging (GO:0019073), viral RNA genome replication (GO:0039694), and viral entry into host cell (GO:0046718) are shown in distinct clusters. In Fig 6 B, the UMAP plot shows the clustering of virus proteins associated with E. coli based on the multi-class model, revealing a greater diversity of viral GO terms, approximately 3.7% (87 out of 2359) were assigned a viral life cycle GO term. In this UMAP plot, ten different GO terms related to the viral life cycle were used to colour the proteins in the UMAP. Viral capsid assembly (GO:0019069), viral entry into host cell (GO:0046718), virion assembly (GO:0019068), viral DNA genome packaging (GO:0019073) and viral DNA genome replication (GO:0039693) are clustered in distinct regions. Within these clusters, there are two sub-clusters viral capsid assembly and virion assembly, which are closely related GO terms. Based on the structure of the GO tree, viral capsid assembly is the child node of virion assembly, indicating that the projection of embeddings with high attention weights is explainable for relationships of GO terms.

The UMAP of the embeddings of the top-ranked viral protein associated with H. sapiens displays a clearer clustering structure (Fig 6 C and D). Overall, the embeddings capture the consistency features of viruses, which are important predictive signals for determining virus-host associations. In Fig 6 C, the UMAP plot is based on the attention weights obtained from the binary model, where 10 different GO terms were retrieved from InterProScan, with nearly 23.9% (1280 out of 5357) of the proteins being assigned GO terms. In UMAP Fig 6 D, UMAP is based on attention weights obtained from a multi-class model, where 10 different GO terms were retrieved from InterProScan, with roughly 12.6% (676 out of 5357) of the proteins being assigned GO terms. We specifically selected protein GO terms related to the viral life cycle for analysis and assigned different colours to each GO term in the UMAP plots Fig 6 C and D. It is worth noting that the number of top 5 ranked proteins based on the multi-class models is fewer than the binary tasks, as more proteins obtained a weight of 0 and were not included in the top-ranked selection.

Again, the UMAP plots demonstrate that different protein functions are clustered into separate locations, while GO terms with similar functions tend to form sub-clusters. This observation highlights the ability of the model to capture and represent the protein function and properties. For example, in both Fig 6 C and D, viral life cycle (GO:0019058), viral RNA genome replication (GO:0039694), and viral genome replication (GO:0019079) occupy a significant percentage of the clusters and are distributed across different projections in the UMAP, where viral life cycle (GO:0019058) shows in diverse clusters as it is the parent node including a wide variety of GO terms. Several embeddings annotated by viral genome replication (GO:0019079) and viral RNA genome replication (GO:0039694) are tightly clustered and close to each other. This observation is consistent with the hierarchical structure of Gene Ontology (GO), where viral RNA genome replication is a subset of viral genome replication. This indicates that ESM-1b has the ability to capture protein functions, which is important for host prediction. UMAPs of binary (Fig 6 C) and multi-class (Fig 6 D) shared the majority of GO terms, except for virion assembly (GO:0019068), which is only presented in UMAP projection of the binary model (Fig 6 C). This suggests that virion assembly (GO:0019068) may play a distinct role in the binary classification task compared to the multi-class classification task.

To visualize the projection of UMAP in proteins, we choose the top 5 ranked proteins to present protein GO annotation clusters. In Fig 7, we can see that the top 5 ranked proteins annotated by GO terms include important proteins with high weights, and proteins from a virus might contain weights of 0, so selecting the top 5 ranked proteins allows us to collect proteins with GO annotations and high weights, as the aim is to present key proteins which are assigned high weights based on binary and multi-class models. We mark the GO annotation for proteins and select two GO annotations as examples. For example, the indices of GO annotations of the top 5 ranked proteins (Fig 7 A) are the same as the indices of all proteins (Fig 7 B), meaning that we can obtain GO annotations although only selecting top 5 ranked proteins.

**Fig 7.**
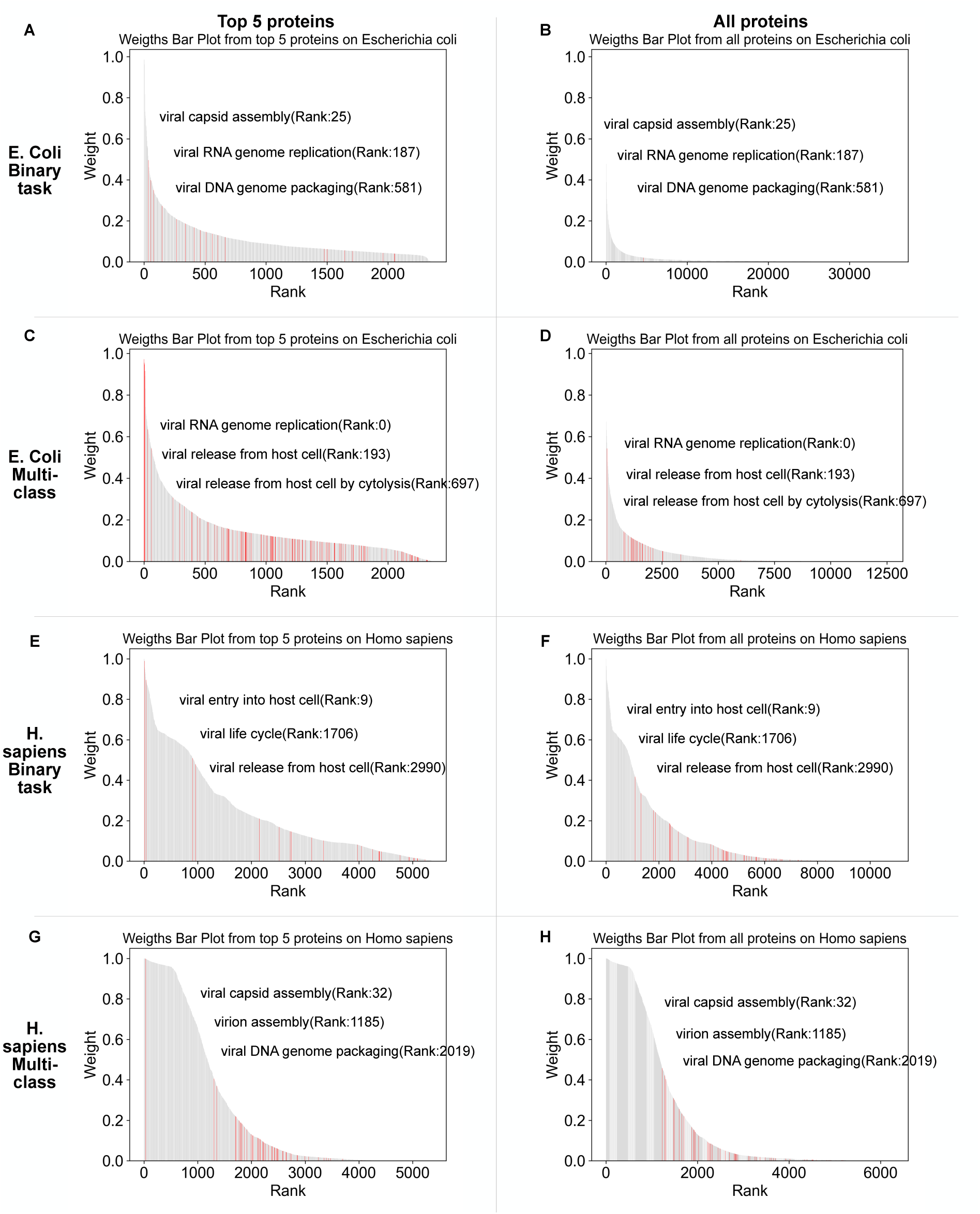
The bar plots display the ranking of weights for the top 5 proteins and all proteins, respectively. The top four bar plots illustrate protein weights obtained for E. coli based on binary classification models(A, B) and multi-class classification models (C, D), respectively. Similarly, the bottom four bar plots depict the protein weights obtained for H. sapiens based on binary classification models(E, F) and multi-class classification models (G, H), respectively. Each host consists of two sections: the left subplot shows the top 5 ranked protein weights, while the right subplot displays all protein weights sorted in descending order.

As shown in Fig 7, this selection allows us to focus on proteins that are assigned high weights, considering the possibility that some proteins from a given virus may have weights of 0. By choosing the top 5 ranked proteins, we are able to gather proteins with GO annotations that also possess high weights. This approach allows us to highlight key proteins assigned high weights based on binary and multi-class models. Fig 7 A and B demonstrate that some ranks of GO annotations for the top 5 ranked proteins aligned with those ranks of all proteins, indicating that GO annotations can still be obtained even when considering only the top 5 ranked proteins.

### 2.5 EvoMIL identifies key proteins of SARS-CoV-2

To highlight EvoMIL performance on an unseen virus, we looked at the prediction and attention weights of SARS-CoV-2 [16] using the trained H. sapiens binary classifier from section 2.2.1. Wuhan-Hu-1 with NCBI Reference Sequence NC 045512.2, which is not included in the training, was predicted to be a human virus with a probability of 0.97 compared to the SVM-kmer binary classifiers [13] of less than 0.01. The top three ranked proteins contributing to host prediction are spike sub-sequences, Non-structural proteins sub-sequences, Nucleocapsid protein, with weights of 0.298, 0.221 and 0.189 respectively. The spike protein contains the receptor binding site which is responsible for cell entry. Nucleocapsid protein is an RNA binding protein that plays a critical role at many stages of the viral life cycle making direct interactions with many host proteins [17]. More N protein and host proteins interactions can be retrieved from protein-protein interaction databases in UniPort (https://www.uniprot.org/uniprotkb/P0DTC9/entry#interaction). The viral process GO terms of these top-ranked proteins are “fusion of virus membrane with the host plasma membrane” (GO:0019064), “fusion of virus membrane with host endosome membrane” (GO:0039654), “receptor-mediated virion attachment to host cell” (GO:0046813) and “endocytosis involved in viral entry into host cell” (GO:0075509). It has been reported that these GO terms are associated with viral infections, indicating that protein attention weights obtained by EvoMIL perform well in identifying important proteins contributing to virus-host associations.

## 3 Discussion

In this paper, we introduce EvoMIL, a novel method for virus-host prediction. Inspired by the success of NLP approaches in biology, we demonstrate the power of using the protein transformer language model ESM-1b to generate embeddings of viral proteins that are highly effective features for virus-host classification tasks. ESM-1b is capturing meaningful biological/host information from viral proteins despite the training dataset being mainly comprised of proteins from cellular life with only 1% being viral proteins. We demonstrate that attention-based MIL can identify which of a virus’s proteins are most important for host prediction, and by implication which proteins may be key to virus-host specificity.

Our classification results show that EvoMIL is able to predict the host at the species level with high AUC, accuracy and F1 scores. The prokaryote binary classifiers achieved a mean AUC of 0.992 whereas the eukaryotic classifiers achieved a mean AUC of 0.851 during selecting negative samples by **Strategy 1**. We also evaluate the model’s performance using different levels of negative sampling from **Strategy 2**, our findings indicate that training binary classifiers becomes more challenging when those hosts associated with negative and positive samples tend to share the same taxonomic levels in the taxonomic tree. Additionally, results demonstrated that eukaryotic host prediction is a more difficult task for two reasons: viruses associated with eukaryotic hosts are much more diverse across all seven Baltimore classes compared to the prokaryotic viruses which are mainly double-stranded DNA viruses; secondly, the eukaryote datasets contain many RNA viruses which have far fewer proteins, this makes it more challenging for MIL which needs many instances in each bag to perform well. Table 1 represents that ESM-1b features outperform more conventional amino acid, physio-chemical properties and DNA k-mer features on multi-class classification tasks. Furthermore, the clustering structure seen in the E. coli and H. sapiens UMAP projections (Fig 6) indicates that embeddings are able to capture functionally related information of the high-ranked proteins associated with host specificity. Moreover, EvoMIL is able to find SARS-cov-2 spike proteins from top-ranked proteins obtained by attention weights of the model.

Transformers enable a one-step pipeline to generate dense feature vectors from viral genomes and learn multiple layers of information about the structural and functional properties of proteins in an unbiased way. Multiple-instance learning is able to capitalize on any host signal contained within individual protein sequences, circumnavigating the need to represent each virus with a single feature vector. This enables it to find common patterns/signals across bags of proteins that differ greatly in both size and content. An additional advantage of attention-based MIL is the ability to identify which proteins are important in prediction, the identification of important proteins of viruses that contribute to host prediction is essential in understanding the specific viral proteins that play a critical role in host infection. Further analysis is needed to test whether this can be exploited to uncover the mechanisms behind virus-host specificity.

In this paper we took a virus-based approach in which only information from the viral genomes is used to generate features, this limits us to classifying viruses for those hosts that have a minimum number of known viruses. Also, by limiting the viral sequences to only Refseq sequences to reduce redundancy, we have further narrowed the range of hosts we can predict. The ultimate host predictor would be able to make predictions for hosts which have no or few known viruses, to do this we will include host information. Combining virus and host-based approaches has been shown to greatly increase the host range of a prediction tool whilst maintaining a low false discovery rate, Roux et al.[18] and Wang et al. [19]. In the future, we will make use of the much larger numbers of host-annotated virus genomes available in public databases and include host information to construct a model that can make predictions for associations for any pair of viruses and host.

## Materials and Methods

### Data

The current study was done using datasets that were collected from the Virus-Host database (VHDB), (https://www.genome.jp/virushostdb/) [11]. The VHDB contains a manually curated set of known species-level virus-host associations collated from a variety of sources, including public databases such as RefSeq, GenBank, UniProt, and ViralZone and evidence from the literature surveys. For each known interaction, this database provides NCBI taxonomic ID for the virus and host species and the Refseq IDs for the virus genomes. We download datasets on 20-10-2021, along with the associated FASTA files containing the raw viral genomes and the FAA file with translated CDS for each viral protein. At this point, the VHDB contained 17733 associations between 12650 viruses and 3740 hosts that were used to construct binary datasets for both prokaryotic and eukaryotic hosts.

### Constructing balanced binary dataset

In this study, we only collect the reference genome from Refseq genome [20] for each virus. The aim is to reduce the amount of redundant or similar sequences in the datasets. We obtain known virus-host associations for prokaryotic and eukaryotic hosts from the Virus-Host database VHDB [11], and remove viruses whose protein sequences do not exist in the Refseq database.

Balanced binary datasets were constructed for each host with an equal number of positive and negative virus sets to obtain a balanced data set for binary classification tasks. For both the prokaryotic and eukaryotic datasets, we collected 4696 associations between 4696 viruses and 498 prokaryotic hosts at the species level; 9595 positive associations from 9595 viruses and 1665 eukaryotic hosts at the species level.

For each binary virus-host association data set, viruses can be represented by *V* = *{V*_1_*, V*_2_*, ..V_V_ },* and the host is represented by *H*. The positive labels consisted of known associations from VHDB [11], and the set of positive viruses can be represented by *V_pos_* = *{V*_1_*, V*_2_*, ..V_P_ }*, where 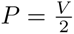. As there is no information on negative associations in the public datasets. Most phages are likely to infect a range of hosts belonging to the same taxonomy [21, 22], therefore, to mitigate possible errors, some researchers [23, 24] construct negative virus-host associations by selecting those viruses from the remaining viruses that do not infect the given host, instead of relating to hosts with different taxa from the given host. using this method, we construct putative negative virus-host associations by collecting viruses from a specific range. As for a given host *H* in the prokaryotic datasets, all viruses are represented by *V_pro_* = *{V*_1_*, V*_2_*, ..V_T_ }*, we collect negative viruses *V_neg_* = *V_pro_ − V_pos_*, which are not related with the given host; and host species taxonomy of *V_neg_* are different from host *H*. On top of that, in order to get the balance datasets for model training and testing, the size of datasets *V_neg_* is equal to *V_pos_*, we randomly choose viruses from putative negative virus set *V_neg_*, finally we get the same number of positive and negative virus datasets, that can be represented by *V_neg_*= *{V_P_* _+1_*, V_P_* _+2_*, ..V_V_ }*. The number of viruses related to the given host is at least 50 and 125 for the prokaryotic and eukaryotic datasets. In addition, there exist some segmented and non-segmented viruses; segmented viruses have multiple RefSeq sequences and need to be combined to represent complete viral sequences. Non-segmented viruses are not divided into multiple segments, it is only necessary to select one RefSeq sequence, although those have multiple virus reference accession numbers. After setting the threshold for the number of viruses at the species level, combining segmented viruses and removing redundancy non-segmented viruses, we have 15 prokaryotic datasets and 5 eukaryotic datasets.

### Feature extraction

The Pre-trained ESM-1b model is used to transform protein sequences into fixed-length embedding vectors that are used as features for downstream tasks such as binary and multi-class classification. For n protein sequences (*h*_1_*, …, h_n_*) input into ESM-1b [8], we obtain n embedding vectors **X** = (**x_1_***, …***x_n_**) each of dimensions 1024. In the pre-training process, the representation is projected to log probabilities (*y*_1_*, …, y_n_*), the model posterior of amino acid at position *i*is represented by a softmax over *y_i_*, the output embeddings (*h*_1_*, …, h_n_*) is applied in downstream tasks, (the parameters are “–repr layers 33 –include mean per tok”). Here, we adapt virus embedding vectors to the supervised Attention-based MIL learning tasks.

There are two special cases of sequences that need to be pre-processed to be suitable as input for ESM-1b:

1. In NCBI, some protein sequences include the amino acid J, which is used to refer to unresolved leucine (L) or isoleucine (I) residues. However, the ESM-1b model does not include the token ’J’ and they must be removed before processing. In order to remove J from NCBI sequences, we randomly replace J with either Leucine(L) or Isoleucine(I).
2. The ESM-1b model can only process sequences with a maximum of 1024 tokens including the beginning-of-sequence (BOS) token and the ending-of-sequence (EOS) token, meaning that the maximum length of the protein sequence is 1022 amino acids. The parameter ’–truncation’ can be used to crop a longer sequence, but this will result in a loss of some part of the sequence information. In order to include information from the entire protein sequence, we split longer sequences and process the sections simultaneously. For protein sequences longer than 1022, we split the sequence into lengths of 1022 amino acids. If the final section is shorter than 25 amino acids it is discarded as it is considered too short to include meaningful information. So that for a given protein sequence *H*, of length *len*(*H*):
  if *len*(*H*) *<* 1022 + 25 then truncate the sequence
  else if *len*(*H*) *≥* 1022 + 25 split the sequence.
  For example, if we have a sequence of length 2049 (2 *×* 1022 + 25), it will be cut into two sub-sequences with the length of 1022, and one sub-sequence with the length of 25. Resulting in three embedding vectors for the single protein sequence that are all assigned to the same label as the parent protein and will be considered instances in the attention-based MIL.

### Attention-based Multiple instance learning

#### Multiple instance learning

We represent a virus as a set of protein embeddings *X* = *{***x**_1_, **x**_2_, **x**_3_*, …,* **x***_M_ }*. The label of the virus is *Y ∈ {*0, 1*}*. If *Y* = 1, the virus is known to be associated with the host, otherwise, the virus is not associated with the host. But individual label *{y*_1_*, y*_2_*, y*_3_*, …, y_M_ }* is unknown. Multiple instance learning (MIL) is used to predict a label for a bag with a set of instances, so it was used to predict a host label for a set of proteins from a virus. The assumption about the label *Y* can be represented as follows:

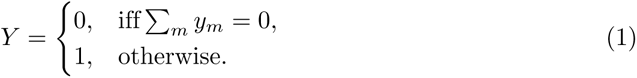

The MIL model can be interpreted as a probabilistic model in which the label of the bag is distributed according to the Bernoulli distribution with the parameter *θ*(*X*) *∈ {*0, 1*}*, which is invariant to the permutation of instances.

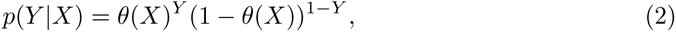

In the multi-class problem, the model can be trained with the softmax loss instead. Given an input instance *x_m_*, the whole scoring function of the MIL can be represented as Eq (3) [25]:

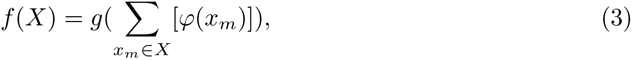

Where, *f* (*X*) is a symmetric function, the orders of the instances do not affect the results of the scoring function *f* (*X*). Here *g* and *φ* are suitable transformations, and the two transformations are modelled using neural networks. Generally, the algorithm can be implemented by modelling a permutation-invariant function *θ*(*X*). (i) *φ*: generate transformed instances; (ii) *f* (*X*): combine transformed instances; (iii) *g*: obtain the final label based on the combined instances.

MIL is the embedding level, instances are mapped to a low-dimensional embedding based on the function *φ*, and then the bag representation can be obtained by MIL pooling. Here the bag representation is not related to the number of instances in the bag, finally, a bag-level classifier was applied to process the bag representation and generate *θ*(*X*).

#### Attention-based pooling

To explain protein embeddings of viruses and learn each feature of protein sequences of viruses, an attention mechanism is considered as MIL pooling [10]. This method applied a weighted average of instances, each instance’s weight is calculated by the neural network, and the sum of weights is 1. Here *U* = *{u*_1_*, u*_2_*, u*_3_*, …, u_M_ }* is a bag of *M* embedding vectors, the MIL pooling is represented by Eq (4):

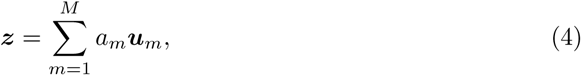

Here, the attention mechanisms can generate a weight for each protein of viruses, and it can be represented as:

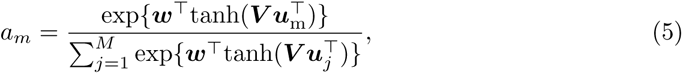

Here, *a_m_* is the weight for the corresponding protein sequence, *w ∈ R^L×^*^1^ and *V ∈ R^L×M^* are parameters, *tanh*(.) is the hyperbolic tangent function [10]. (Code is available from https://github.com/AMLab-Amsterdam/AttentionDeepMIL?utm_source=catalyzex.com). We obtained these weights to quantify the importance of proteins to host prediction.

### Evaluation

To evaluate our classifiers we use six evaluation metrics, including AUC, accuracy, F1 score, sensitivity, specificity and precision as defined below. The two main evaluation metrics used in our analysis and explanation are AUC and accuracy. The area under the receiver operating characteristic curve (AUC) is used to evaluate machine learning performance; accuracy is the ratio of the number of correctly predicted samples to the total number of samples.

True positive (TP), true negative (TN), false positive (FP), and false negative (FN) are the parameters used to calculate specificity, sensitivity, and accuracy. True positive(TP): predicted label and known label are positive. True Negative(TN): predicted label and known label are negative. False Positive (FP): The predicted label is positive, but the actual label is negative. False Negative(FN): The predicted label is negative, but the actual label is positive. Sensitivity is the percentage of positive samples; specificity is the percentage of negative samples. The F1 score is the harmonic mean of precision and recall. Here, we set average=’macro’ to calculate the F1 score and precision for each label and get their unweighted mean. The formula of the evaluation indices is as follows:

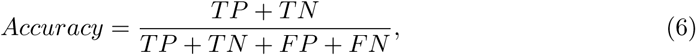

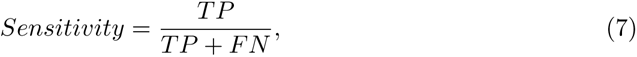

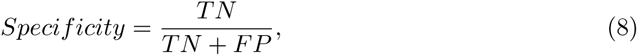

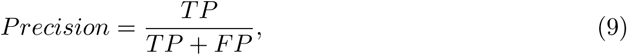

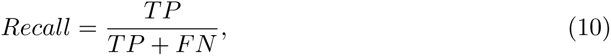

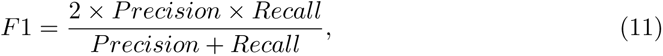

## Acknowledgments

This research was funded by European Union’s Horizon 2020 research and innovation program, under the Marie Sk-lodowska-Curie Actions Innovative Training Networks grant agreement no. 955974 (VIROINF). The authors also acknowledge funding from the Medical Research Council (MRC, MC UU 12014/12), the Biotechnology and Biological Sciences Research Council (BBSRC, BB/V016067/1), Engineering and Physical Sciences Research Council (EPSRC, EP/R018634/1) and the European Union’s Horizon 2020 research and innovation programme project PANCAIM (101016851). The authors thank Kieran Lamb for the helpful discussion.

## Appendix S1

Code and trained models are available on https://github.com/liudan111/EvoMIL.

**Table S1. Statistic table of 22 prokaryotic hosts.** The table of 22 prokaryotic datasets shows the host species name, the number of viruses associated with each host, and the average, minimal and maximal number of protein sequences of viruses associated with each host.

**Table S2. Statistic table of 36 eukaryotic hosts.** The table of 36 eukaryotic datasets shows the host species name, the number of viruses associated with each host, and the average, minimal and maximal number of protein sequences of viruses associated with each host.

**Table S3. Results of binary classifiers in prokaryotic hosts.** Evaluation indices are obtained by testing 5-fold cross-validation models on each host, and then the mean and standard deviation of the evaluation metrics can be obtained. Evaluation metrics include AUC, accuracy, f1, specificity, sensitivity, and precision.

**Table S4. Results of binary classifiers in eukaryotic hosts.** Evaluation indices are obtained by testing 5-fold cross-validation models on each host, and then the mean and standard deviation of the evaluation metrics can be obtained. Evaluation metrics include AUC, accuracy, f1, specificity, sensitivity, and precision.

**Table S5. Table of accuracy based on ESM-1b and K-mers.** The accuracy are calculated for prokaryotic and eukaryotic hosts associated with fewer viruses, ranging from 5 to 30.

**Table S6. Table of GO ID and GO terms.** To understand the protein functions from viruses, we retrieve the GO term for each GO ID, which is presented in the UMAP plots.

**Fig. S1 The Confusion matrix plot of prokaryotic hosts based on EvoMIL.** The confusion matrix plot represents the performance of the EvoMIL model on 22 prokaryotic hosts. It is constructed by evaluating the model’s predictions on a test set comprising 20% of the dataset, while the EvoMIL model was trained on the remaining 80% of the data. This plot provides insights into the model’s accuracy in predicting the host species for the tested viruses.

**Fig. S2 The Confusion matrix plot of eukaryotic hosts based on EvoMIL.** The confusion matrix plot represents the performance of the EvoMIL model on 36 eukaryotic hosts. It is constructed by evaluating the model’s predictions on a test set comprising 20% of the dataset, while the EvoMIL model was trained on the remaining 80% of the data. This plot provides insights into the model’s accuracy in predicting the host species for the tested viruses.

---

**Table S1.** Prokaryotic hosts: hostname, the number of viruses associated with the host. The table of 22 prokaryotic datasets shown the host species name, the number of viruses associated with each host, and the average, minimal and maximal number of protein sequences of viruses associated with each host.

| Index | Host name | Number of viruses | Mean_protein | Min_protein | Max_protein |
| --- | --- | --- | --- | --- | --- |
| 1 | Mycolicibacterium smegmatis | 838 | 105 | 58 | 258 |
| 2 | Escherichia coli | 474 | 134 | 4 | 611 |
| 3 | Salmonella enterica | 231 | 117 | 15 | 276 |
| 4 | Lactococcus lactis | 185 | 56 | 22 | 194 |
| 5 | Pseudomonas aeruginosa | 182 | 91 | 3 | 375 |
| 6 | Gordonia terrae | 176 | 96 | 57 | 233 |
| 7 | Staphylococcus aureus | 152 | 86 | 18 | 233 |
| 8 | Klebsiella pneumoniae | 121 | 95 | 26 | 299 |
| 9 | Cutibacterium acnes | 81 | 45 | 42 | 48 |
| 10 | Enterococcus faecalis | 80 | 85 | 23 | 224 |
| 11 | Bacillus thuringiensis | 62 | 163 | 30 | 304 |
| 12 | Acinetobacter baumannii | 60 | 106 | 29 | 326 |
| 13 | Vibrio cholerae | 59 | 77 | 6 | 230 |
| 14 | Arthrobacter sp. ATCC 21022 | 55 | 76 | 26 | 114 |
| 15 | Bacillus cereus | 54 | 127 | 25 | 292 |
| 16 | Microbacterium foliorum | 39 | 60 | 25 | 155 |
| 17 | Erwinia amylovora | 39 | 212 | 49 | 345 |
| 18 | Cellulophaga baltica | 36 | 87 | 13 | 198 |
| 19 | Ralstonia solanacearum | 35 | 56 | 3 | 343 |
| 20 | Synechococcus sp. WH 7803 | 33 | 200 | 51 | 292 |
| 21 | Listeria monocytogenes | 32 | 122 | 38 | 196 |
| 22 | Paenibacillus larvae | 31 | 72 | 56 | 92 |

**Fig S1.**
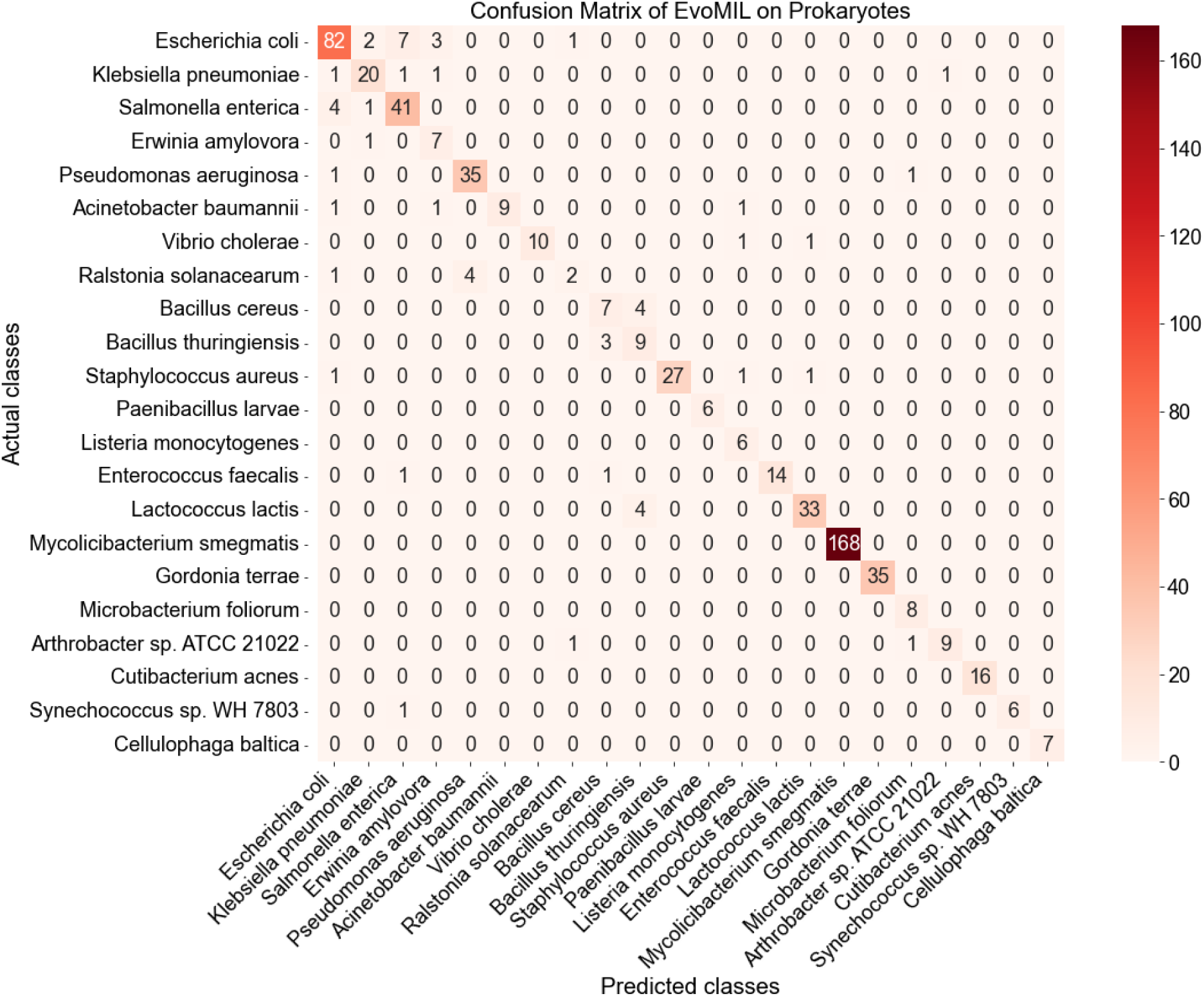
The Confusion matrix plot of prokaryotic hosts based on EvoMIL. The confusion matrix plot represents the performance of the EvoMIL model on 22 prokaryotic hosts. It is constructed by evaluating the model’s predictions on a test set comprising 20% of the dataset, while the EvoMIL model was trained on the remaining 80% of the data. This plot provides insights into the model’s accuracy in predicting the host species for the tested viruses.

**Fig S2.**
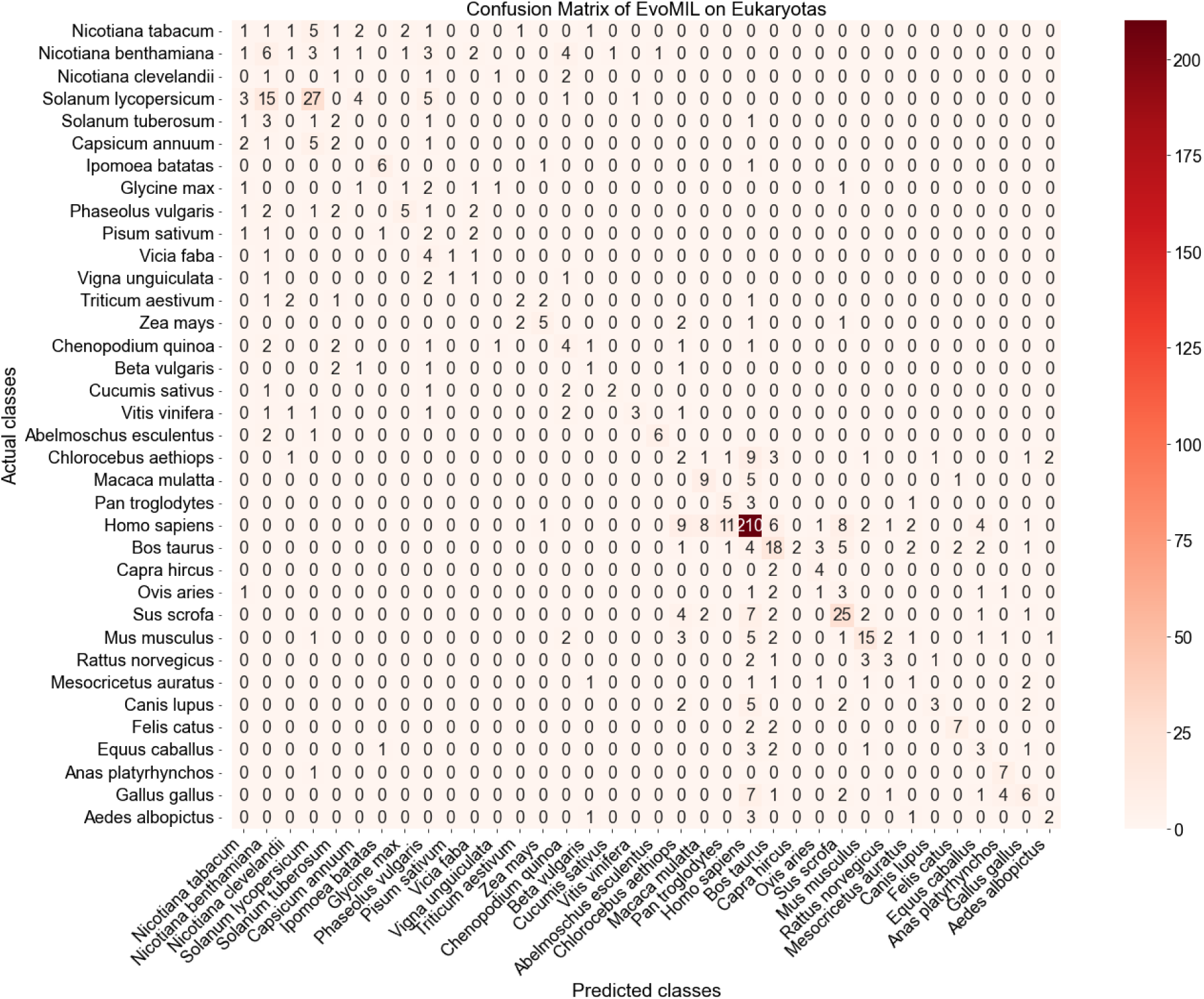
The Confusion matrix plot of eukaryotic hosts based on EvoMIL. The confusion matrix plot represents the performance of the EvoMIL model on 36 eukaryotic hosts. It is constructed by evaluating the model’s predictions on a test set comprising 20% of the dataset, while the EvoMIL model was trained on the remaining 80% of the data. This plot provides insights into the model’s accuracy in predicting the host species for the tested viruses.

**Table S2.** Eukaryotic hosts: hostname and the number of viruses associated with the host. The table of 36 eukaryotic datasets shown the host species name, the number of viruses associated with each host, and the average, minimal and maximal number of protein sequences of viruses associated with each host.

| Index | Host name | Number of viruses | Mean_protein | Min_protein | Max_protein |
| --- | --- | --- | --- | --- | --- |
| 1 | Homo sapiens | 1321 | 8 | 1 | 262 |
| 2 | Solanum lycopersicum | 277 | 4 | 1 | 13 |
| 3 | Sus scrofa | 219 | 10 | 1 | 184 |
| 4 | Bos taurus | 202 | 14 | 1 | 236 |
| 5 | Mus musculus | 174 | 18 | 1 | 233 |
| 6 | Nicotiana benthamiana | 123 | 4 | 1 | 13 |
| 7 | Gallus gallus | 109 | 15 | 1 | 261 |
| 8 | Chlorocebus aethiops | 108 | 23 | 1 | 220 |
| 9 | Nicotiana tabacum | 81 | 4 | 1 | 13 |
| 10 | Macaca mulatta | 73 | 18 | 1 | 223 |
| 11 | Canis lupus | 72 | 10 | 1 | 233 |
| 12 | Phaseolus vulgaris | 69 | 4 | 1 | 13 |
| 13 | Chenopodium quinoa | 66 | 3 | 1 | 13 |
| 14 | Felis catus | 57 | 12 | 1 | 233 |
| 15 | Capsicum annuum | 55 | 3 | 1 | 8 |
| 16 | Equus caballus | 54 | 22 | 1 | 236 |
| 17 | Zea mays | 53 | 5 | 1 | 14 |
| 18 | Ovis aries | 51 | 13 | 1 | 150 |
| 19 | Vitis vinifera | 49 | 4 | 1 | 12 |
| 20 | Rattus norvegicus | 48 | 12 | 1 | 233 |
| 21 | Abelmoschus esculentus | 47 | 3 | 1 | 9 |
| 22 | Solanum tuberosum | 46 | 3 | 1 | 10 |
| 23 | Pan troglodytes | 45 | 10 | 1 | 169 |
| 24 | Triticum aestivum | 43 | 5 | 1 | 14 |
| 25 | Ipomoea batatas | 42 | 5 | 1 | 11 |
| 26 | Mesocricetus auratus | 42 | 5 | 1 | 11 |
| 27 | Glycine max | 41 | 4 | 1 | 14 |
| 28 | Anas platyrhynchos | 40 | 3 | 1 | 29 |
| 29 | Aedes albopictus | 36 | 2 | 1 | 11 |
| 30 | Vicia faba | 34 | 4 | 1 | 9 |
| 31 | Pisum sativum | 33 | 3 | 1 | 8 |
| 32 | Nicotiana clevelandii | 32 | 4 | 1 | 13 |
| 33 | Cucumis sativus | 32 | 4 | 1 | 12 |
| 34 | Vigna unguiculata | 32 | 4 | 1 | 13 |
| 35 | Beta vulgaris | 32 | 5 | 1 | 10 |
| 36 | Capra hircus | 31 | 16 | 1 | 150 |

**Table S3.** Results for prokaryotic hosts. **Results of binary classifiers in prokaryotic hosts.** Evaluation indices are obtained by testing 5-fold cross-validation models on each host, then the mean and standard deviation of each evaluation metric can be obtained. Evaluation metrics include AUC, accuracy, f1, specificity, sensitivity, and precision.

| Host name | AUC | Accuracy | f1 | specificity | sensitivity | precision |
| --- | --- | --- | --- | --- | --- | --- |
| Mycobacterium smegmatis | 0.999±0.0 | 0.993±0.0 | 0.993±0.0 | 0.998±0.0 | 0.99±0.0 | 0.993±0.0 |
| Escherichia coli | 0.947±0.01 | 0.893±0.02 | 0.892±0.02 | 0.868±0.07 | 0.915±0.03 | 0.896±0.02 |
| Salmonella enterica | 0.965±0.03 | 0.899±0.02 | 0.897±0.02 | 0.94±0.02 | 0.851±0.06 | 0.904±0.01 |
| Lactococcus lactis | 0.999±0.0 | 0.995±0.01 | 0.995±0.01 | 1.0±0.0 | 0.988±0.02 | 0.995±0.01 |
| Pseudomonas aeruginosa | 0.978±0.01 | 0.953±0.01 | 0.953±0.01 | 0.956±0.02 | 0.951±0.02 | 0.954±0.01 |
| Gordonia terrae | 0.998±0.0 | 0.975±0.01 | 0.975±0.01 | 0.966±0.01 | 0.983±0.02 | 0.975±0.01 |
| Staphylococcus aureus | 0.974±0.01 | 0.964±0.01 | 0.963±0.01 | 0.962±0.0 | 0.966±0.02 | 0.963±0.02 |
| Klebsiella pneumoniae | 0.986±0.02 | 0.963±0.03 | 0.963±0.03 | 0.978±0.05 | 0.945±0.04 | 0.966±0.03 |
| Cutibacterium acnes | 0.68±0.44 | 0.842±0.22 | 0.751±0.34 | 0.6±0.55 | 1.0±0.0 | 0.721±0.38 |
| Enterococcus faecalis | 0.995±0.0 | 0.944±0.01 | 0.934±0.02 | 1.0±0.0 | 0.922±0.02 | 0.917±0.02 |
| Bacillus thuringiensis | 0.992±0.01 | 0.928±0.02 | 0.927±0.02 | 1.0±0.0 | 0.88±0.03 | 0.924±0.02 |
| Acinetobacter baumannii | 0.984±0.02 | 0.933±0.02 | 0.929±0.02 | 1.0±0.0 | 0.84±0.05 | 0.949±0.02 |
| Vibrio cholerae | 0.957±0.03 | 0.925±0.03 | 0.924±0.04 | 0.969±0.04 | 0.873±0.05 | 0.931±0.04 |
| Arthrobacter sp. ATCC 21022 | 0.953±0.02 | 0.873±0.05 | 0.873±0.05 | 0.873±0.08 | 0.873±0.05 | 0.875±0.05 |
| Bacillus cereus | 0.977±0.03 | 0.864±0.03 | 0.862±0.03 | 0.982±0.04 | 0.745±0.04 | 0.886±0.03 |

**Table S4.** Results for eukaryotic hosts. **Results of binary classifiers in eukaryotic hosts.** Evaluation indices are obtained by testing 5-fold cross-validation models on each host, and then the mean and standard deviation of each evaluation metric can be obtained. Evaluation metrics include AUC, accuracy, f1, specificity, sensitivity, and precision.

| Host name | AUC | Accuracy | f1 | specificity | sensitivity | precision |
| --- | --- | --- | --- | --- | --- | --- |
| Homo sapiens | 0.918±0.01 | 0.84±0.01 | 0.84±0.01 | 0.856±0.03 | 0.823±0.04 | 0.842±0.01 |
| Solanum lycopersicum | 0.851±0.01 | 0.8±0.03 | 0.8±0.03 | 0.808±0.06 | 0.793±0.03 | 0.801±0.03 |
| Sus scrofa | 0.761±0.01 | 0.727±0.02 | 0.726±0.02 | 0.791±0.04 | 0.667±0.04 | 0.733±0.02 |
| Bos taurus | 0.898±0.0 | 0.837±0.02 | 0.835±0.02 | 0.895±0.03 | 0.774±0.07 | 0.844±0.01 |
| Mus musculus | 0.762±0.02 | 0.697±0.01 | 0.696±0.01 | 0.703±0.02 | 0.691±0.03 | 0.697±0.01 |

**Table S5.** The accuracy (%) of multi-class MIL by using ESM-1b and k-mer features on tiny host datasets. Comparison of mean accuracy and standard deviation between ESM-1b and k-mer feature sets: ESM-1b, AA 2, PC 4 and DNA 5. For each feature, training multi-class classification models on 22 prokaryotic hosts and 36 eukaryotic hosts based on 5-fold cross-validation, then mean and standard deviation of accuracy is obtained by testing the trained model on host datasets that are associated with fewer viruses, ranging from 5 to 30 (tiny host datasets).

| Host type | Features | Genus | Family | Order | Class | Phylum | Kingdom |
| --- | --- | --- | --- | --- | --- | --- | --- |
| Prokaryotes | ESM-1b | **1.8±0.332** | **4.386±0.446** | **9.412±0.61** | **22.928±1.61** | **31.872±0.939** | **97.41±0.0** |
|  | AA_2 | 1.088±0.529 | 5.216±0.894 | 12.22±2.223 | 29.334±2.671 | 39.122±3.93 | 97.41±0.0 |
|  | PC_3 | 1.178±0.258 | 4.556±1.063 | 11.12±2.445 | 25.646±2.7 | 32.868±4.513 | 97.41±0.0 |
|  | DNA_5 | 1.23±0.326 | 4.028±0.479 | 9.542±1.067 | 21.308±1.665 | 25.644±2.253 | 97.41±0.0 |
| Eukaryotes | ESM-1b | **0.908±0.143** | **3.842±0.628** | **5.498±0.589** | **36.868±1.54** | **39.992±1.446** | **47.416±1.41** |
|  | AA_2 | 0.328±0.142 | 2.086±0.225 | 4.8±0.31 | 24.006±1.761 | 29.844±1.248 | 43.222±0.555 |
|  | PC_3 | 0.334±0.063 | 2.19±0.174 | 4.826±0.602 | 27.374±1.492 | 32.374±1.372 | 44.344±0.915 |
|  | DNA_5 | 0.42±0.178 | 2.102±0.319 | 4.366±0.644 | 25.144±3.519 | 31.792±2.37 | 43.908±1.225 |

**Table S6.** Table of GO ID and GO terms. The GO terms are coloured in the UMAP plots, we retrieve protein functions for each GO ID, and GO terms are shown in the table.

| GO ID | GO terms |
| --- | --- |
| GO:0019076 | viral release from host cell |
| GO:0046718 | viral entry into host cell |
| GO:0044659 | viral release from host cell by cytolysis |
| GO:0039693 | viral DNA genome replication |
| GO:0019069 | viral capsid assembly |
| GO:0019068 | virion assembly |
| GO:0019073 | viral DNA genome packaging |
| GO:0039694 | viral RNA genome replication |
| GO:0019058 | viral life cycle |
| GO:0019062, GO:0046718 | virion attachment to host cell, viral entry into host cell |
| GO:0019062 | virion attachment to host cell |
| GO:0019079 | viral genome replication |
| GO:0046813 | receptor-mediated virion attachment to host cell |
| GO:0019072 | viral genome packaging |
| GO:0019076, GO:0046718 | viral release from host cell, viral entry into host cell |

